# Semi-Automated Detection, Annotation, and Prognostic Assessment of Ictal Chirps in Intracranial EEG from Patients with Epilepsy

**DOI:** 10.1101/2025.08.13.670167

**Authors:** Nooshin Bahador, Milad Lankarany

## Abstract

We analyzed the spectro-temporal characteristics of ictal “chirp” events in intracranial EEG (iEEG) recordings from 13 epilepsy patients, using a custom derivative dataset of 22,721 spectrograms that we generated from the Epilepsy-iEEG-Multicenter Dataset. Ictal chirps, transient frequency-modulated patterns, were semi-automatically annotated to assess their relationship with seizure onset zones (SOZs) and surgical outcomes. Preprocessing included notch filtering (60 Hz, 120 Hz) and bandpass filtering (1–60 Hz), followed by segmentation into 60-second windows. Spectrograms were generated via Short-Time Fourier Transform (STFT) with a Hann window (87.5% overlap) and converted to dB scale. Chirps were annotated by manually drawing bounding boxes, followed by automated ridge detection, model fitting, and feature extraction (start/end time-frequency, duration, direction, RMSE, R^2^). Spatial, Spectro-temporal, and clinical features were analyzed using heatmaps, hierarchical clustering, statistical tests (Mann-Whitney U), and outcome prediction models. Patient-channel mappings revealed clustering of chirp patterns among specific patient pairs, correlating with shared clinical profiles. Flow-based analysis demonstrated prognostic value: very high spectral durations in SOZ regions were associated with favorable surgical outcomes (80.43% success rate), whereas very high temporal durations in SOZ correlated with poorer outcomes (51.35% risk). Statistical comparisons showed significant differences between SOZ and non-SOZ chirps: SOZ chirps exhibited longer spectral durations (10.13 ± 6.35 Hz vs. 8.51 ± 5.66 Hz, *p* < 0.001), shorter temporal durations (6.76 ± 5.83 s vs. 7.14 ± 5.39 s, *p* = 0.006), and higher spectro-temporal ratios (2.66 vs. 1.92, *p* < 0.001). Distribution analyses further indicated that prolonged temporal chirps were more prevalent in non-SOZ regions.

## 1. Introduction

Seizures are characterized by abrupt and temporary bursts of abnormal electrical activity in the brain, typically resulting from an imbalance between excitatory and inhibitory signals. These events can lower neuronal firing thresholds and disrupt normal brain function. When such seizures occur repeatedly without provocation, the condition is defined as epilepsy, a chronic neurological disorder with a wide spectrum of clinical presentations. Causes of seizures are multifactorial, ranging from hypoxia and genetic mutations to developmental anomalies and medical conditions. The manifestations vary, affecting motor control, awareness, and perception, which complicates both diagnosis and therapeutic targeting (Freeman et al., 1993).

Electroencephalography (EEG) is a key tool in identifying epileptiform activity and classifying it into ictal (during a seizure), inter-ictal (between seizures), and post-ictal (after a seizure) phases. The ictal phase encompasses initiation, propagation, and termination, with many studies focusing on the mechanisms that drive these transitions (e.g., Miri et al., 2018; Rich et al., 2020). Evidence indicates that seizure initiation often involves synchronized inhibition, leading to sparse neuronal spiking and elevated extracellular potassium levels (de Curtis and Avoli, 2016). These dynamic processes can give rise to distinct time-frequency patterns, including “chirps”—gradual frequency increases or decreases within a narrow frequency band (Grinenko et al., 2018).

In epilepsy research, chirp-like features have been observed in intracranial EEG recordings, particularly during seizures (Bahador et al., 2024; Bahador et al., 2025; Li et al., 2020; Gnatkovsky et al., 2011; Kurbatova et al., 2016; Sen et al., 2007; Niederhauser et al., 2003; Schiff et al., 2000; Feltane et al., 2013; Gnatkovsky et al., 2019a; Benedetto and Colella, 1995). Clinical findings suggest that these fast-frequency chirps may serve as reliable indicators of epileptogenic zones (Di Giacomo et al., 2024), although discrepancies exist in when these chirps emerge—ranging from before seizure onset to during ictal propagation (e.g., Kurbatova et al., 2016; Niederhauser et al., 2003).

The researchers reported that the presence and removal of chirp-generating regions correlated with favorable surgical outcomes, highlighting chirps as valuable markers for guiding treatment. Di Giacomo et al. (2024) found that fast activity “chirp” patterns in SEEG recordings are reliable indicators of the epileptogenic zone (EZ) in patients with focal drug-resistant epilepsy. These chirps, marked by shifting high-frequency signals, were consistently detected across diverse seizure types and brain regions. Their presence often matched the areas identified through clinical evaluation, and their removal was associated with better surgical outcomes. The study supports using chirp detection as a practical, objective tool to enhance EZ localization, particularly in complex or MRI-negative cases (Di Giacomo et al., 2024).

Although recent studies have highlighted the potential of ictal chirps as biomarkers for seizure onset zones (SOZs) and predictors of surgical outcomes, a comprehensive characterization of these patterns—spanning their spatiotemporal features and clinical relevance—remains limited. In this study, we utilized the Epilepsy-iEEG-Multicenter Dataset (Li et al., 2023) to systematically analyze chirp dynamics using a multi-modal analytical framework. We created a derivative dataset comprising 22,721 annotated spectrograms, generated through a combination of semi-automated chirp detection and advanced signal processing techniques. Our approach included: (1) spatial mapping of chirps and SOZs via patient-channel heatmaps; (2) clinical phenotype clustering using geometric metrics derived from Engel and ILAE outcome scores (Wieser et al., 2001); (3) predictive modeling of surgical outcomes based on chirp duration and SOZ involvement; and (4) comparative statistical analysis of spectral and temporal chirp features across SOZ and non-SOZ regions.

## 2. Methodology

The Epilepsy-iEEG-Multicenter-Dataset on OpenNeuro includes high-resolution intracranial EEG (iEEG) data from 13 patients across four centers (excluding Cleveland Clinic due to data sharing restrictions). Each patient’s dataset contains raw iEEG recordings organized in BIDS format, with associated metadata including clinical information (Li et al., 2022). A derivative dataset was generated in this work from the original Epilepsy-iEEG-Multicenter-Dataset (hosted on the OpenNeuro repository), consists of approximately 22,721 spectrograms. The presence of chirp patterns, along with their characteristics, was semi-automatically annotated in this dataset. We’ve made the derivative dataset and supplementary materials publicly available on **GitHub**. The spectrogram generation process consists of 6 steps, illustrated in **Figure 1**. The pipeline begins with signal preprocessing, where notch filters remove power line noise at 60 Hz and 120 Hz, and a bandpass filter isolates frequencies between 1 and 60 Hz. Next, the continuous signal is segmented into 60-second windows, ensuring manageable analysis blocks. The Short-Time Fourier Transform (STFT) is then applied using a Hann window with 87.5% overlap to compute the power spectrum over time. The spectral data are converted to the decibel (dB) scale for better interpretability, and frequency content is restricted to 0–60 Hz to focus on relevant neural bands. Finally, the processed data are visualized as a spectrogram, with color intensity representing power in dB, and key parameters such as frequency and time resolution are derived based on STFT settings.

**Figure 1.**
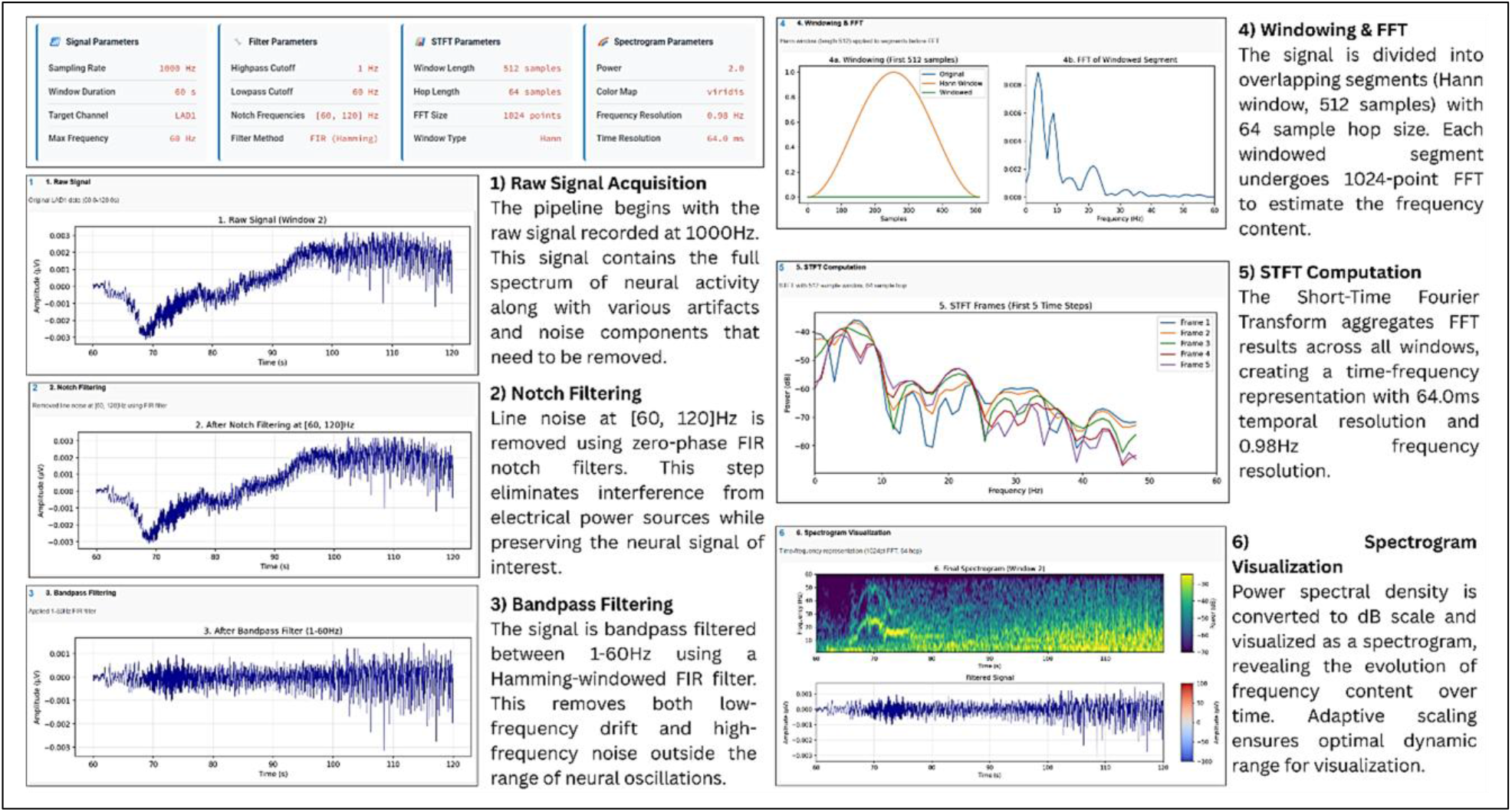
Flowchart illustrating the spectrogram generation pipeline. The process begins with signal preprocessing, including notch and bandpass filtering, followed by segmentation into fixed-length windows. Short-Time Fourier Transform (STFT) is applied using a Hann window with high overlap, and the resulting power spectra are converted to decibel (dB) scale. Frequency content is then limited to a maximum of 60 Hz, and the final output is visualized as a color-coded spectrogram.

Once 2D spectrogram representations were generated and stored, a semi-automated annotation (illustrated in **Figure 2**), combining manual user input with automated processing, was used to create labeled spectrogram datasets. Users manually drew bounding boxes around regions of interest, while the system automatically performed ridge detection, fitted appropriate models, calculated quality metrics (RMSE, R^2^), and determined directionality. An example of manual bounding box annotation combined with automated ridge detection and model fitting is shown in **Figure 3**. The resulting dataset included precise time-frequency coordinates, model parameters, and visualization outputs stored in structured formats (Excel/CSV). This hybrid approach combines human expertise in pattern recognition with algorithmic precision in measurement and modeling.

**Figure 2.**
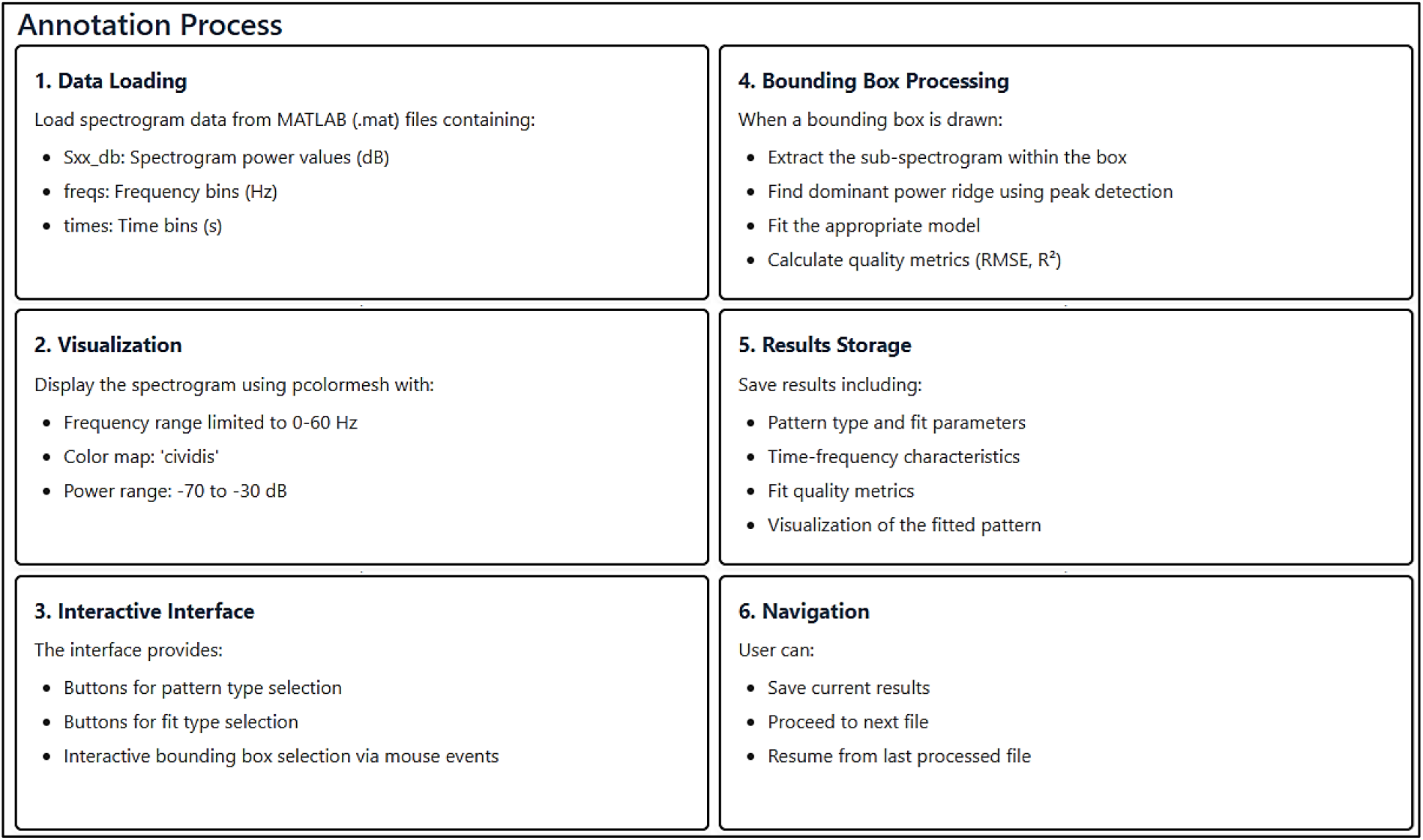
Process of interactive annotation for detecting and characterizing two chirp patterns in spectrograms: The annotation process begins with the user selecting a pattern type and drawing a bounding box around the pattern of interest using mouse events. Within this bounding box, the algorithm identifies the dominant power ridge, to which the code fits either linear or exponential models (see **Figure 3**). The fitting process involves multiple strategies with different initial parameters, includes a constraint to pass through the starting point (initial mouse click), calculates RMSE and R^2^ for assessing fit quality, and determines the direction of the ridge (upward or downward). The results are stored in a pandas Data Frame and saved to an Excel file, including the pattern type, fit parameters, time-frequency coordinates, duration, frequency span, fit quality metrics, and visualization images.

**Figure 3.**
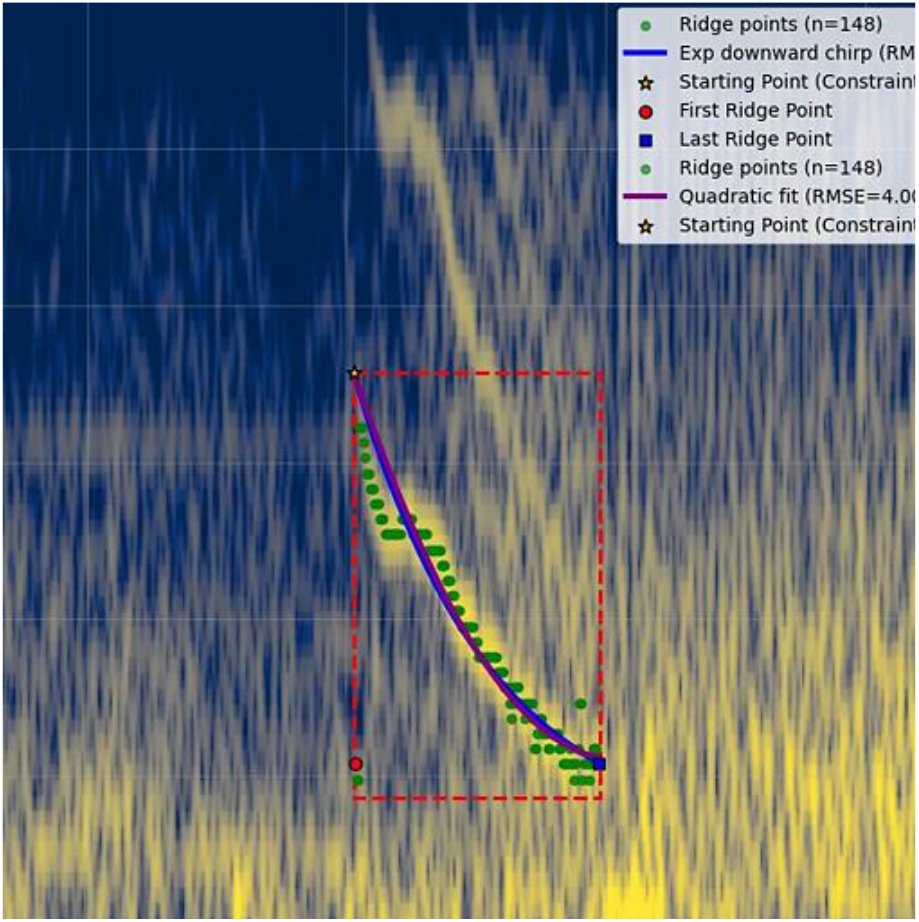
Manual Bounding Box Annotation Combined with Automated Ridge Detection and Model Fitting (Sample spectrogram from patient jh102, run-05, channel-RPF4, window-2)

The analytical approach used in this work is explained in **Figure 4**. We combined spatial, temporal, and clinical data analyses to investigate ictal chirp events in intracranial EEG (iEEG) recordings. The features can be grouped into spatial (e.g., electrode position), spectro-temporal, and clinical categories. Spatial features include: channel (EEG channel location), issoz (seizure onset zone label), isbadcontact (bad contact flag), and ezhypobrainregion (hypothesized epileptogenic brain region). Spectro-temporal features include: starttime (chirp start time), endtime (chirp end time), startfreq (chirp starting frequency), endfreq (chirp ending frequency), durationtime (chirp time duration), durationfreq (chirp frequency range), fitparams (chirp fitting parameters), direction (chirp fitting direction), rmse (chirp fit error - root mean square error), rsquared (chirp fit quality - R^2^ value). Clinical features consist of: clinicaldifficulty (case difficulty), engelscore (post-surgical outcome score), ilaescore (ILAE outcome score), outcome (surgical result). The methodology included: (1) patient-channel heatmap visualizations with optimized sorting (patients are sorted by patient ID, and channels are sorted alphabetically) and scaling to examine spatial distributions of chirp and seizure onset zone (SOZ) events; (2) clinical profile clustering using geometric shape metrics derived from normalized clinical variables (Clinical variables including clinical_difficulty, engel_score, and ilae_score were z-score normalized using StandardScaler to ensure comparability across metrics) with hierarchical clustering; (3) outcome prediction analysis through a three-stage flow process examining SOZ status, duration characteristics, and Engel outcomes; and (4) comparative statistical analyses (Mann-Whitney U tests) of spectral/temporal features between SOZ and non-SOZ regions, supplemented by duration binning analyses and effect size calculations. The effect size is calculated to quantify the magnitude of differences between groups using the Mann-Whitney U test, which compares the distributions of two independent samples. The effect size is computed as 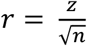, where *z* is the z-score derived from the p-value using the formula 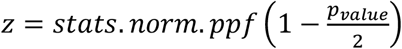 and *n* is the total sample size. The resulting effect size *r* indicates the strength of the difference between groups. The multi-modal approach integrated visualization techniques (heatmaps, radar plots, dendrograms, flow diagrams) with quantitative statistical methods to characterize chirp events and their clinical correlations.

**Figure 4.**
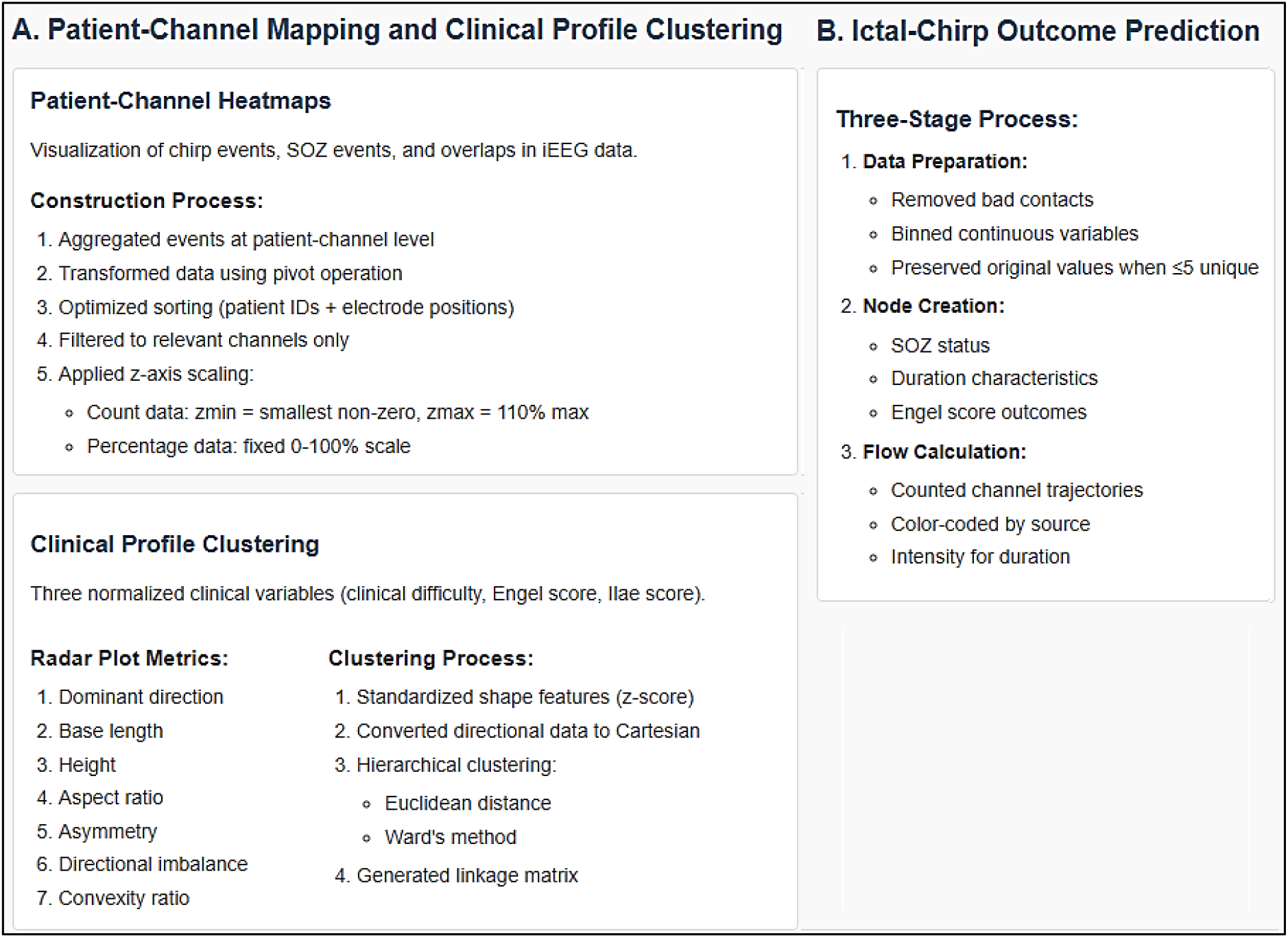
Analytical Framework for Ictal-Chirp Characterization. The figure summarizes the multi-modal methodology integrating: (i) *spatial mapping* via patient-channel heatmaps (lexicographical/electrode-sorted, z-scaled); (ii) *clinical clustering* of normalized profiles (Engel/ILAE scores) using geometric shape metrics and Ward’s hierarchical clustering; (iii) *outcome prediction* through SOZ-duration-outcome flow analysis with trajectory quantification; and (iv) *statistical comparisons* of spectral/temporal features (Mann-Whitney U tests, duration binning, effect sizes). Visualizations (heatmaps, radar plots, dendrograms) are paired with quantitative analyses to correlate chirp properties with clinical outcomes.

## 3. Results

Given semi-automatic chirp detection, we, here, investigated different ictal-chirp characteristics as well as their relationship with patients’ outcomes.

### 3.1. Patient-Channel Mapping and Clinical Profile Clustering of Chirp Events

To visualize the spatial distribution of chirp events, seizure onset zone (SOZ) events, and their overlaps in intracranial EEG (iEEG) data analysis, patient-channel heatmaps were created. Events were first aggregated at the patient-channel level, capturing the frequency of occurrences per recording site. The aggregated data were then transformed using a pivot operation. As shown in **Figure 5**, heatmaps were optimized through a combination of lexicographical sorting of patient identifiers and electrode position-based sorting of channels. Additionally, the data were filtered to include only those channels containing relevant (ictal-chirp) events. Z-axis scaling was applied based on the type of data. For count data, the minimum z-axis value (*zmin*) was set to the smallest non-zero value observed, while the maximum (*zmax*) was defined as 110% of the maximum value to prevent saturation. For percentage data, a fixed scale ranging from 0% to 100% was used to maintain consistency across visualizations.

**Figure 5.**
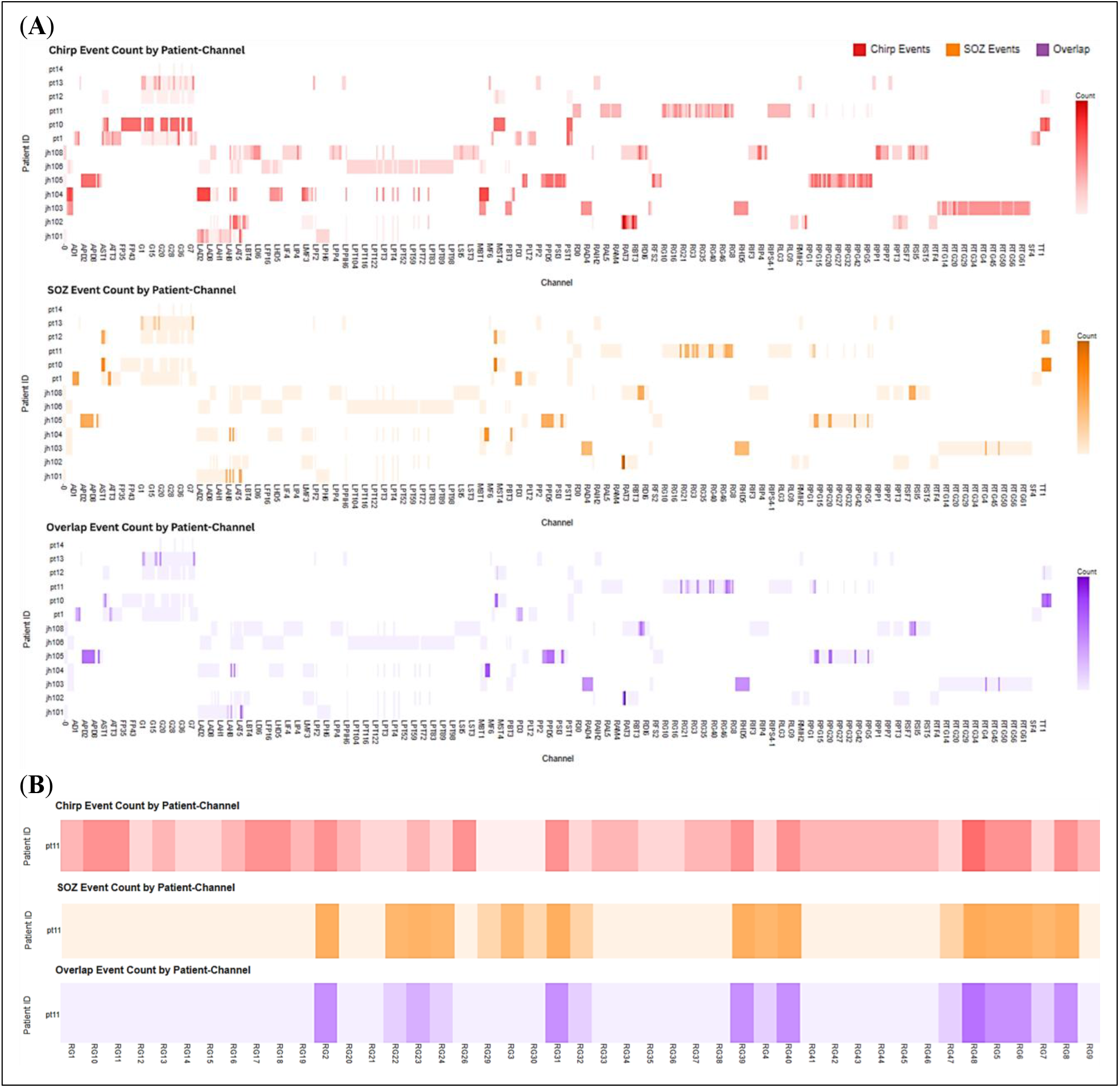
Chirp Detection and Mapping: Patient-Channel Distribution of Chirp and SOZ Events. The heatmap illustrates event counts across patients and channels, with color intensity indicating event frequency with different scales. (**A**) Top panel: Distribution of chirp events by patient and channel. Middle panel: Distribution of SOZ events by patient and channel. Bottom panel: Overlap of chirp and SOZ. (**B**) Zoomed-in view of patient PT11, focusing on the channels from the RG group.

For each patient data, we created a clinical profile including three key numeric clinical variables, namely, clinical difficulty, Engel score, and Ilae score, were selected and normalized to a [0,1] range to allow for comparative analysis across different scales. Each patient’s clinical profile was represented as a triangle, shown by radar plot in **Figure 6**, using these normalized variables, from which seven geometric shape metrics were computed: dominant direction (circular mean of angles), base length (longest side), height (from opposite vertex to base), aspect ratio (height/base), asymmetry (deviation from an equilateral triangle), directional imbalance (vector sum magnitude), and convexity ratio (hull area to polygon area). Each triangle in the radar plot represents a single patient’s clinical profile, based on three normalized variables: clinical difficulty, Engel score, and ILAE score (scaled to [0,1]). The shape of each triangle varies depending on the patient’s unique clinical parameters. Some triangles are large and expansive, while others are small, compact, elongated, or asymmetric. These geometric differences reflect variations in clinical complexity across the three dimensions. Each triangle serves as a “*clinical fingerprint*,” and from these shapes, we extract seven geometric metrics, including dominant direction, base length, height, and aspect ratio—to quantitatively describe each patient. The goal of this analysis is to investigate whether patients with similar clinical fingerprints also exhibit similar chirp distributions across their recorded channels (see **Figure 7**). All shape features, excluding the directional component, were standardized using z-score normalization, while the directional data was converted to Cartesian coordinates via trigonometric transformation for compatibility with Euclidean distance metrics. Hierarchical clustering was then performed using Euclidean distance as the similarity measure and Ward’s minimum variance method to form compact, variance-minimizing clusters. This process generated a linkage matrix capturing the hierarchical structure of cluster merges, including child nodes, inter-cluster distances, and cluster sizes at each step.

**Figure 6.**
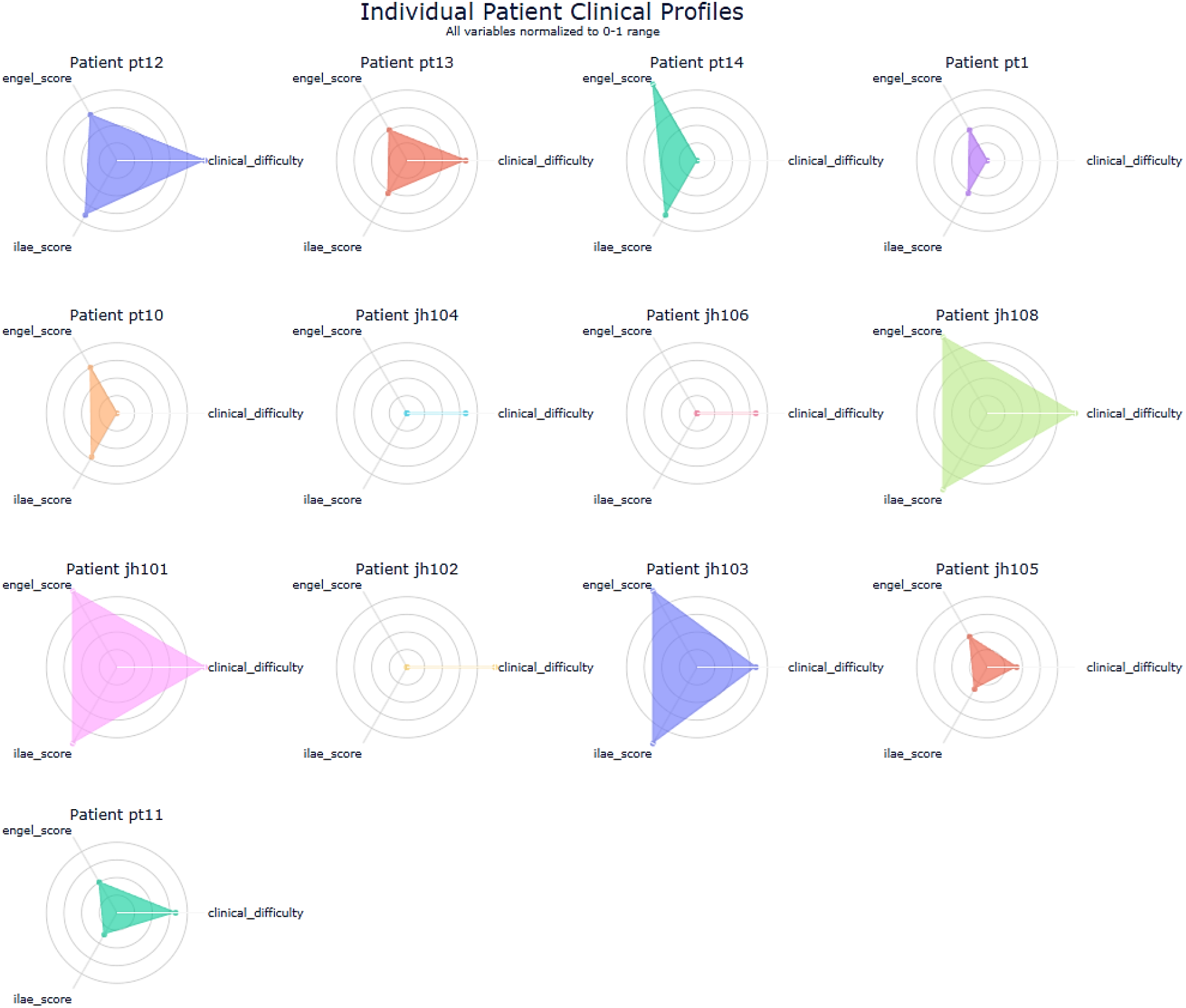
Radar Plot Representation of Clinical Profiles. Patient profiles were shown using geometric shapes in a radar plot. Each triangle represents one patient’s clinical data from three normalized variables: clinical difficulty, Engel score, and ILAE score (all scaled 0–1). Triangle shapes vary—large, small, elongated, or asymmetric—reflecting differences in clinical complexity. These triangles act as “clinical fingerprints.” From each, we extract seven geometric metrics like dominant direction, base length, height, and aspect ratio to describe patients quantitatively. The analysis explores if patients with similar fingerprints have similar chirp distributions across channels.

**Figure 7.**
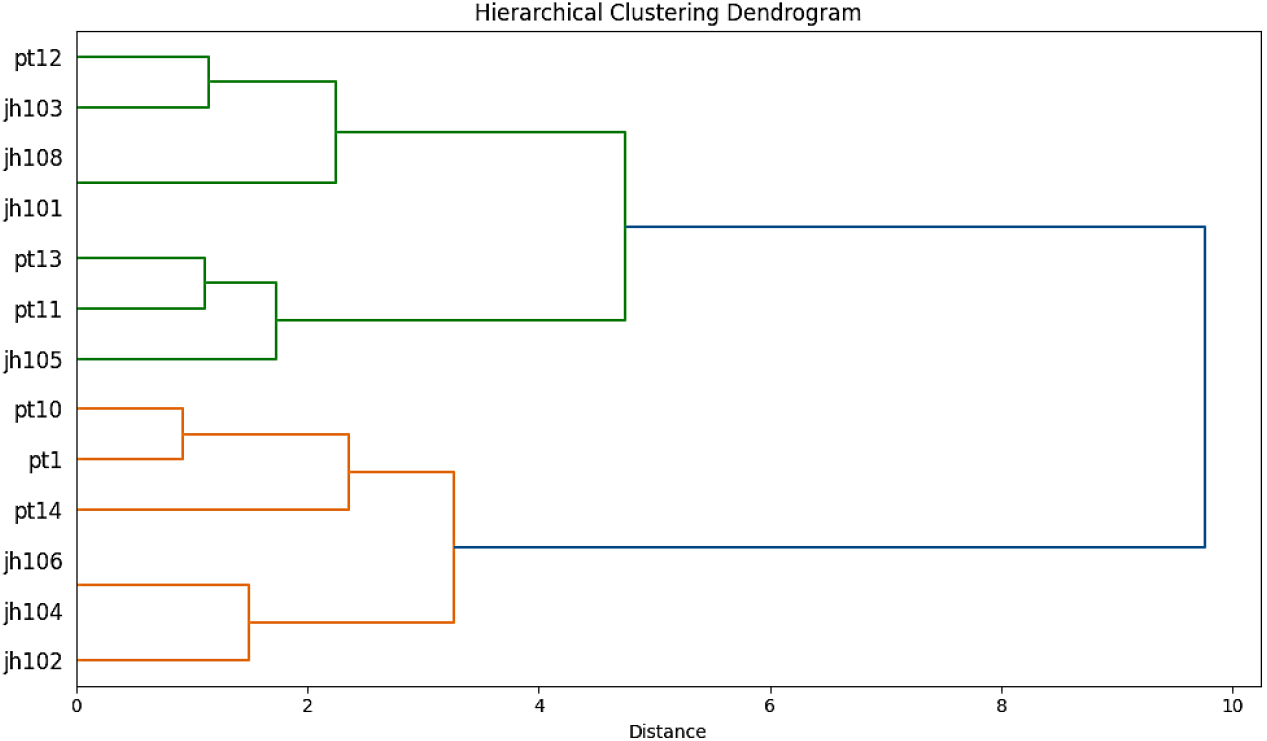
Hierarchical Clustering Dendrogram of Clinical Profiles. Dendrogram showing hierarchical clustering results based on Euclidean distances between standardized shape features (with directional data transformed to Cartesian coordinates). Clustering was performed using Ward’s minimum variance method. Leaf nodes correspond to individual patients, while branch lengths indicate the distance between merged clusters, reflecting similarity among clinical shape profiles.

One can visualize patients’ clinical clustering in **Figure 7**, a right-oriented dendrogram with leaf nodes representing individual patients, branch lengths reflecting inter-cluster distances, and the x- and y-axes denoting merge distances and patient IDs, respectively. This figure presents patient groupings derived from clinical similarity, visualized through a dendrogram. Patients are clustered based on the resemblance of their geometric clinical profiles, with branch lengths representing the degree of similarity, shorter branches indicate more closely related profiles. This clustering approach facilitates the identification of distinct clinical subtypes and provides a framework for exploring whether patients with similar clinical characteristics also share comparable chirp distribution patterns across recorded channels.

### 3.2. Relationship between Ictal-Chirp Characteristics and Patient Outcome

To assess whether Ictal-chirp characteristics can be used as predictors for patients outcomes, we investigated the relationship between Ictal-chirp temporal and spectral features, and patients’ outcomes. This mapping is shown in **Figure 8** and **9** using a three-stage process, namely, **Data Preparation:** Raw clinical data was cleaned by removing bad contacts (is_bad_contact = True). Continuous variables (duration_time, duration_freq) were binned into quartiles when sufficient unique values existed, preserving the original values when ≤5 unique values were present. **Node Creation:** Three node layers were considered: (1) SOZ status (SOZ/Non-SOZ). (2) Duration characteristics represented in four ranges, namely, low, medium, high, very high. The Ictal-chirp duration features were shown in Figure **8** and **9** for temporal and spectral components, respectively. (3) Engel score outcomes. Nodes were sized proportionally to the number of channels in each category. **Flow Calculation:** Directed flows between nodes were calculated by counting channel trajectories through each path. Flows were colored by source category (red for SOZ, blue for Non-SOZ) with intensity indicating duration characteristics (teal/yellow/purple/orange for low/medium/high/very-high). According to these diagrams, a surgeons could potentially use these ictal-chirp duration profiles (temporal and spectral) during pre-surgical evaluation to better predict which patients are likely to benefit from epilepsy surgery.

**Figure 8.**
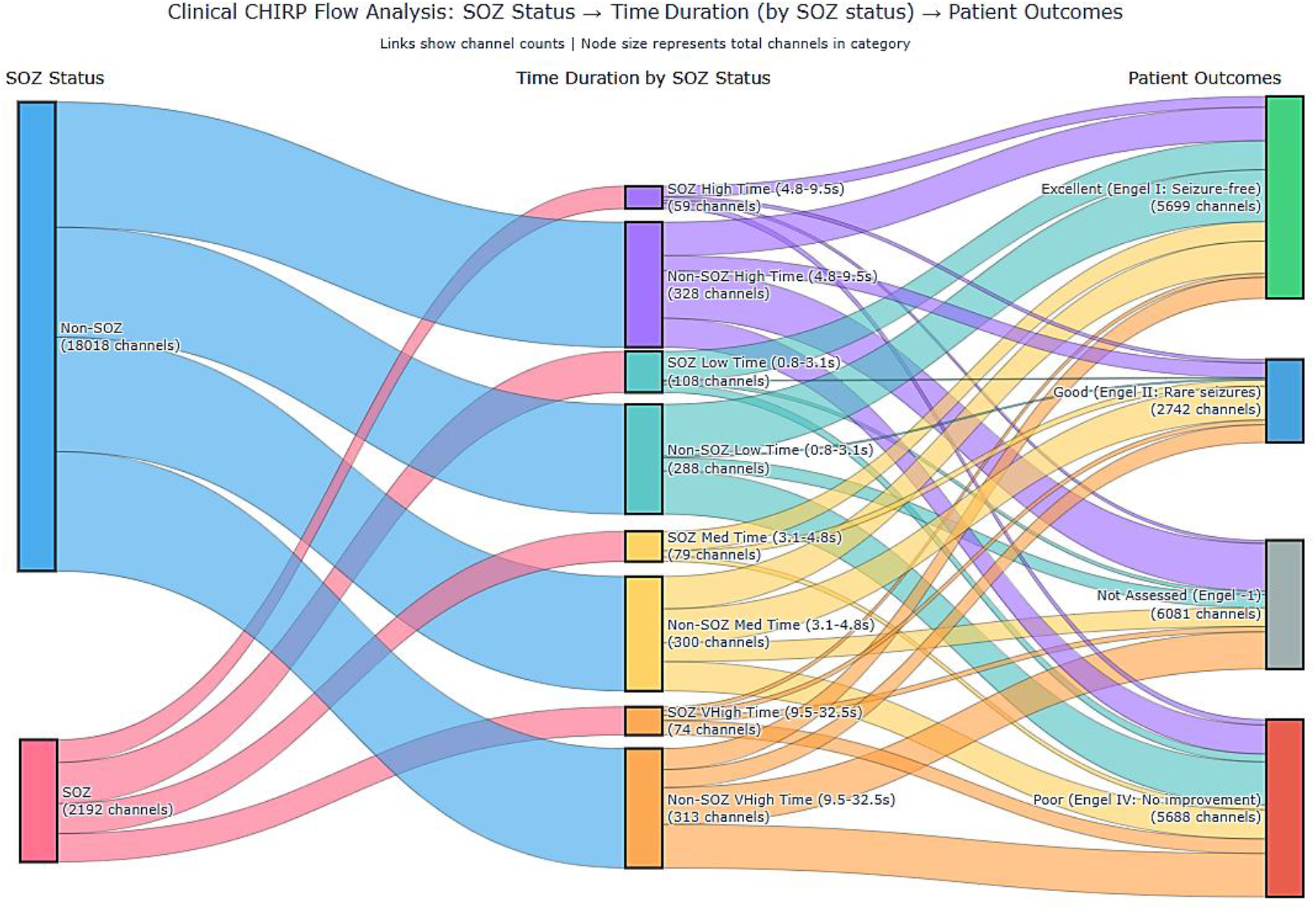
Temporal Duration Analysis: This diagram shows the flow: SOZ Status → Time Duration Categories (separated by SOZ status) → Patient Outcomes. Time duration categories are based on quartiles of the actual data distribution, with separate nodes for SOZ and Non-SOZ channels.

**Figure 9.**
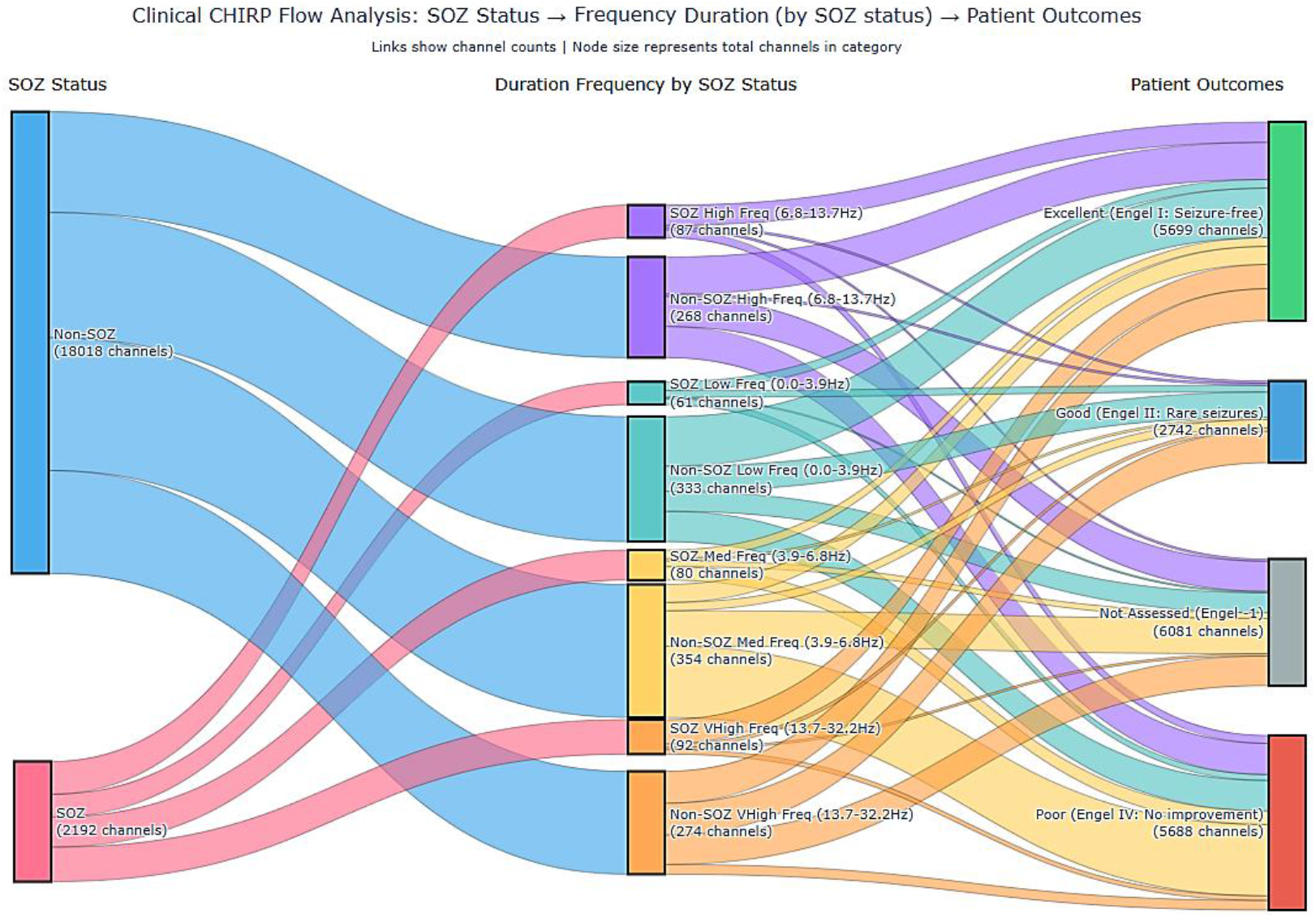
Spectral Duration Analysis: This diagram shows the flow: SOZ Status → Duration Frequency Categories (separated by SOZ status) → Patient Outcomes. Frequency Duration categories are based on quartiles of the actual data distribution, with separate nodes for SOZ and Non-SOZ channels.

One can observe that the highest risk bins for temporal duration are the Low bin for Non-SOZ (Risk: 39.24%, Count: 113/288) and the Very High (VHigh) bin for SOZ (Risk: 51.35%, Count: 38/74), while for spectral duration, the highest risk bins are the Medium (Med) bin for both Non-SOZ (Risk: 53.67%, Count: 190/354) and SOZ (Risk: 45.00%, Count: 36/80). The most favorable bins for good or excellent outcomes based on temporal duration are the Medium (Med) bin for both Non-SOZ (58.00%, Count: 174/300) and SOZ (84.81%, Count: 67/79), while for spectral duration, the most favorable bins are the Very High (VHigh) bin for both Non-SOZ (61.68%, Count: 169/274) and SOZ (80.43%, Count: 74/92) (Refer to the detailed outcome data provided in Appendix I.).

### 3.3. Spatio-temporal features of Ictal Chirp

Ictal chirps contain both spatial and temporal information, each playing a role in understanding the seizure onset zone (SOZ). **Figure 10** shows representative CHIRP patterns in spectrograms from sub jh102 (run-05, channel RPF4, window II), jh103 (run-02, MBT3, window I), pt1 (run-03, PST4, window II), and jh105 (run-05, APD7, window I). The pie chart in **Figure 11** visualizes the relative proportion of events originating from seizure onset zones versus non-seizure onset zones.

**Figure 10.**
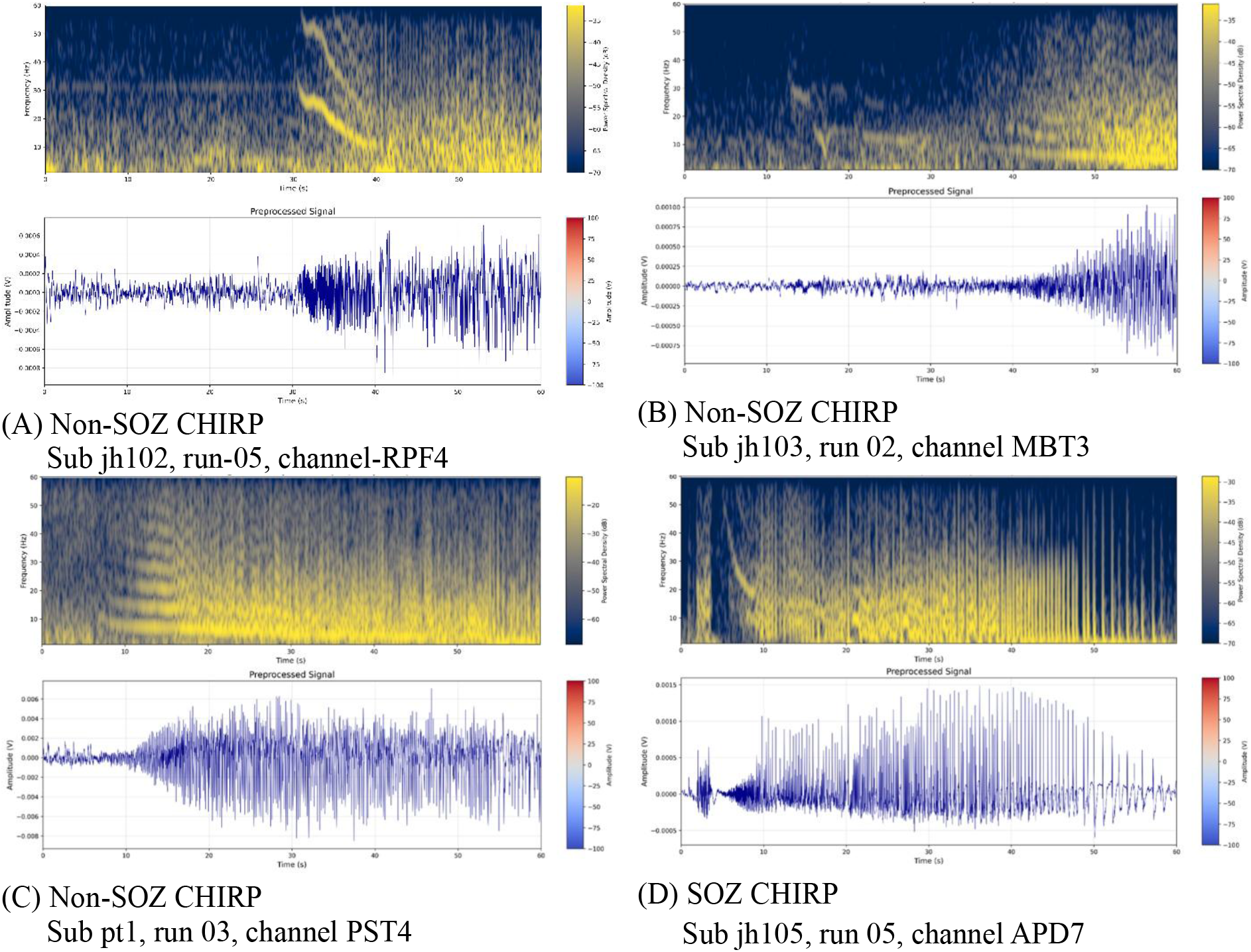
Representative CHIRP Patterns Visible in Spectrograms

**Figure 11.**
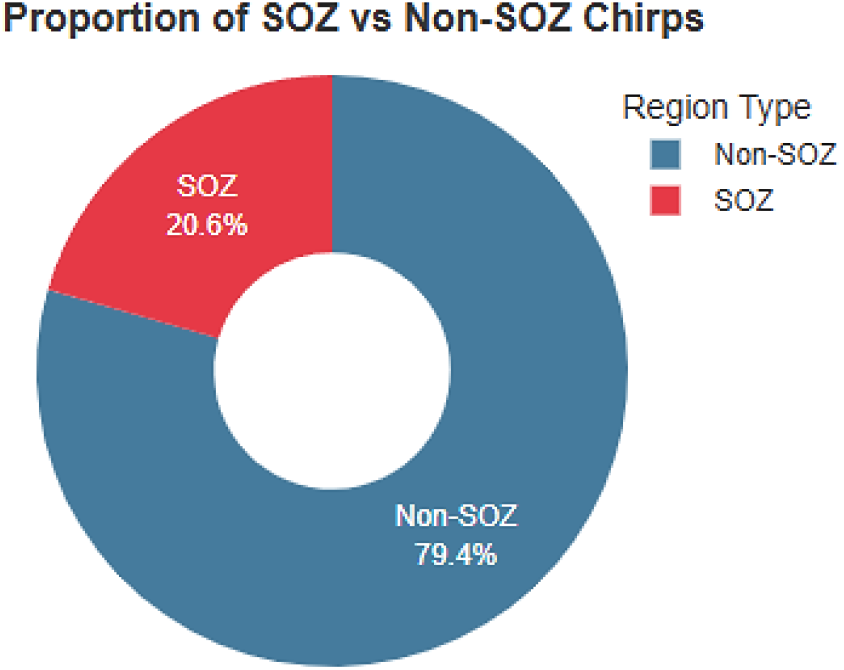
Distribution of CHIRP events between SOZ and Non-SOZ regions: The data contains 1546 CHIRPs after filtering, with 318 (20.6%) originating from SOZ regions and 1228 (79.4%) from Non-SOZ regions.

To compare distribution of spectro-temporal durations between SOZ and non-SOZ Chirps, we draw Box plots in **Figure 12**. **Figure 12** (**A**) compares spectral duration distributions between SOZ and Non-SOZ CHIRPs. The box shows quartiles (25th to 75th percentile), the line represents the median, and whiskers extend to 1.5× the interquartile range. Points beyond this are considered outliers. Statistical annotation shows Mann-Whitney U test results. Violin plots in **Figure 12** (**B**) comparing time duration distributions. The width represents kernel density estimation of the probability density, showing the full distribution shape. Statistical annotation shows test results. Box plots in **Figure 12** (**C**) compare the ratio of frequency to time duration. This metric may reveal differences in how frequency evolves over time between SOZ and Non-SOZ regions. The statistical analysis of CHIRP characteristics in **Table 1** reveals significant differences between seizure onset zone (SOZ) and non-seizure onset zone (Non-SOZ) regions across measured metrics. Specifically, the spectral duration of CHIRP events was significantly longer in SOZ areas (10.13 ± 6.35 Hz) compared to Non-SOZ regions (8.51 ± 5.66 Hz), with a small effect size (r = 0.10, p = 0.000). Conversely, the temporal duration was slightly shorter in SOZ (6.76 ± 5.83 s) than in Non-SOZ (7.14 ± 5.39 s), though this difference was still statistically significant (p = 0.006, r = 0.07). The spectro-temporal ratio was notably higher in SOZ (2.66 ± 2.18) compared to Non-SOZ (1.92 ± 1.74), showing the strongest effect among the three measures (r = 0.12, p = 0.000). Despite all differences reaching statistical significance, the small effect sizes suggest that while these CHIRP features differ between SOZ and Non-SOZ regions, although the magnitude of these differences might be modest.

**Figure 12.**
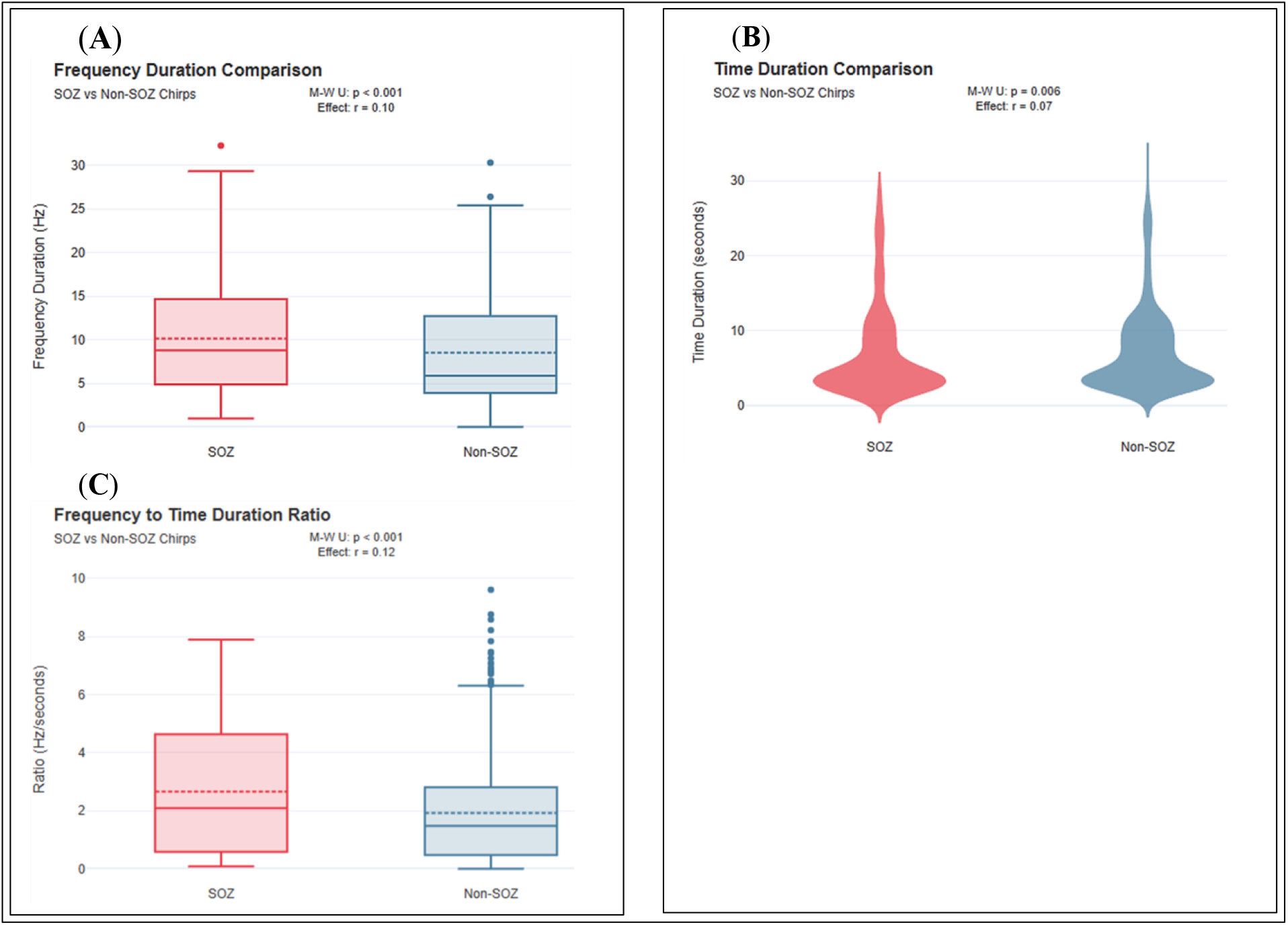
**(A)** Spectral Duration Comparison of CHIRPs. **(B)** Temporal Duration Comparison of CHIRPs. **(C)** Spectro-Temporal Ratio Comparison of CHIRPs.

**Table 1.** Statistical Analysis: Frequency duration of CHIRP: Significantly different between SOZ and Non-SOZ (p = 0.000, effect size r = 0.10). Time duration of CHIRP: Significantly different between SOZ and Non-SOZ (p = 0.006, effect size r = 0.07). Spectro-Temporal Ratio of CHIRP: Significantly different between SOZ and Non-SOZ (p = 0.000, effect size r = 0.12). Statistical tests performed using Mann-Whitney U test with effect size calculation (r = z/√n).

| Metric | SOZ<br>(Mean ± SD) | Non-SOZ<br>(Mean ± SD) | Statistical<br>Significance | Effect<br>Size |
| --- | --- | --- | --- | --- |
| Spectral Duration<br>(Hz) | 10.13 ± 6.35 | 8.51 ± 5.66 | **p = 0.000** | $r = 0.10$ |
| Temporal Duration<br>(s) | 6.76 ± 5.83 | 7.14 ± 5.39 | **p = 0.006** | $r = 0.07$ |
| Spectro-Temporal<br>Ratio | 2.66 ± 2.18 | 1.92 ± 1.74 | **p = 0.000** | $r = 0.12$ |

To further investigate the difference between distribution of SOZ and non-SOZ chirps, we plotted their temporal and spectral (duration) distributions in bar charts with more precise time-frequency scales. Grouped bar chart in **Figure 13** shows the distribution of SOZ and Non-SOZ CHIRPs across equal-frequency bins of frequency duration. Bins were created using quantile binning to ensure equal sample sizes in each bin. This reveals whether certain frequency ranges of CHIRPs are more associated with SOZ regions. Grouped bar chart in **Figure 14** shows the distribution across time duration bins.

**Figure 13.**
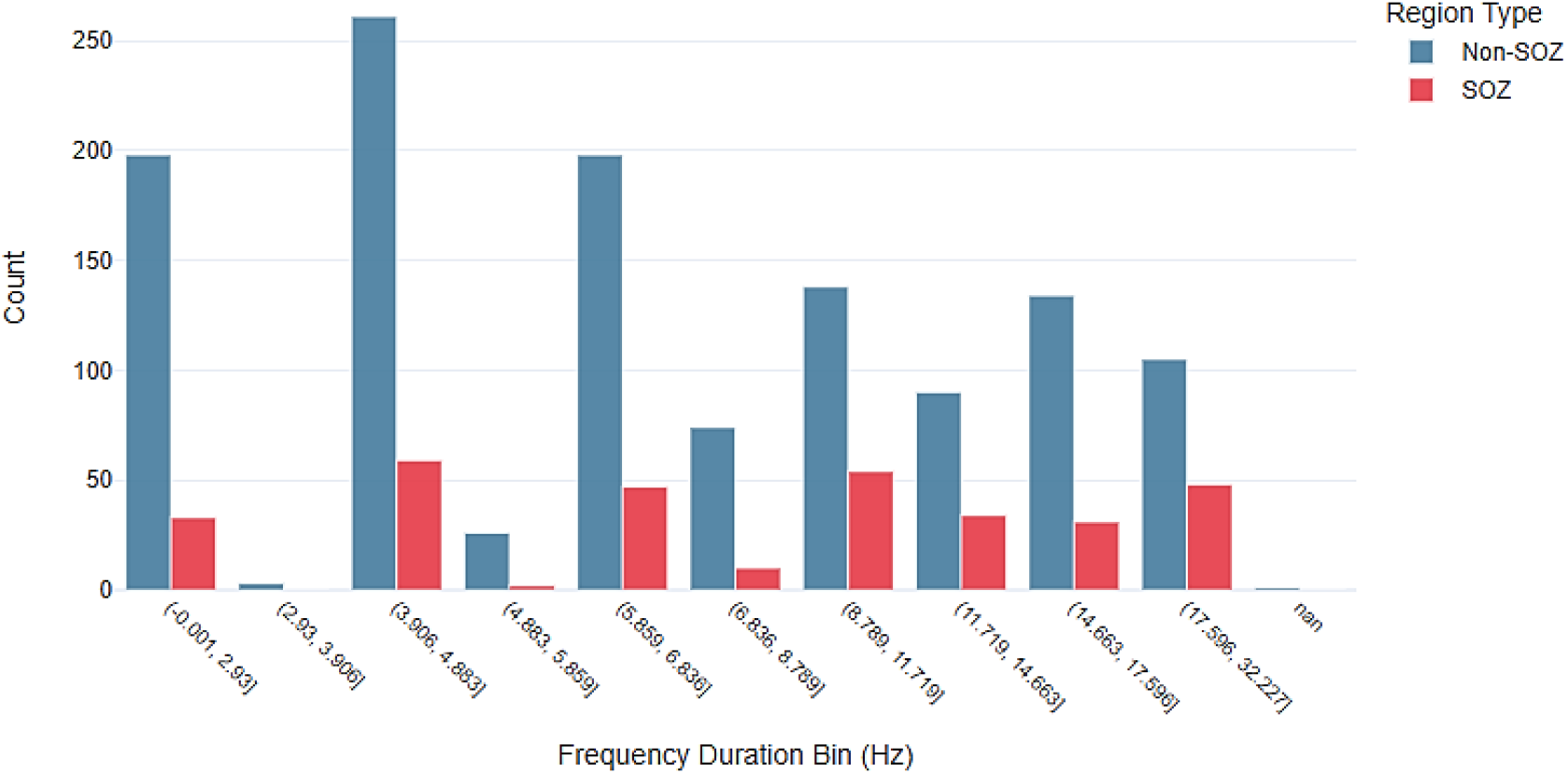
Distribution Across Spectral Duration Bins of CHIRPs: There is a general downward trend in the latter half of the chart, indicating that Non-SOZ CHIRPs are more frequent in the lower spectral duration bins. As spectral duration increases, the proportion of Non-SOZ CHIRPs declines, suggesting that these regions tend to produce CHIRPs with shorter spectral spans.

**Figure 14.**
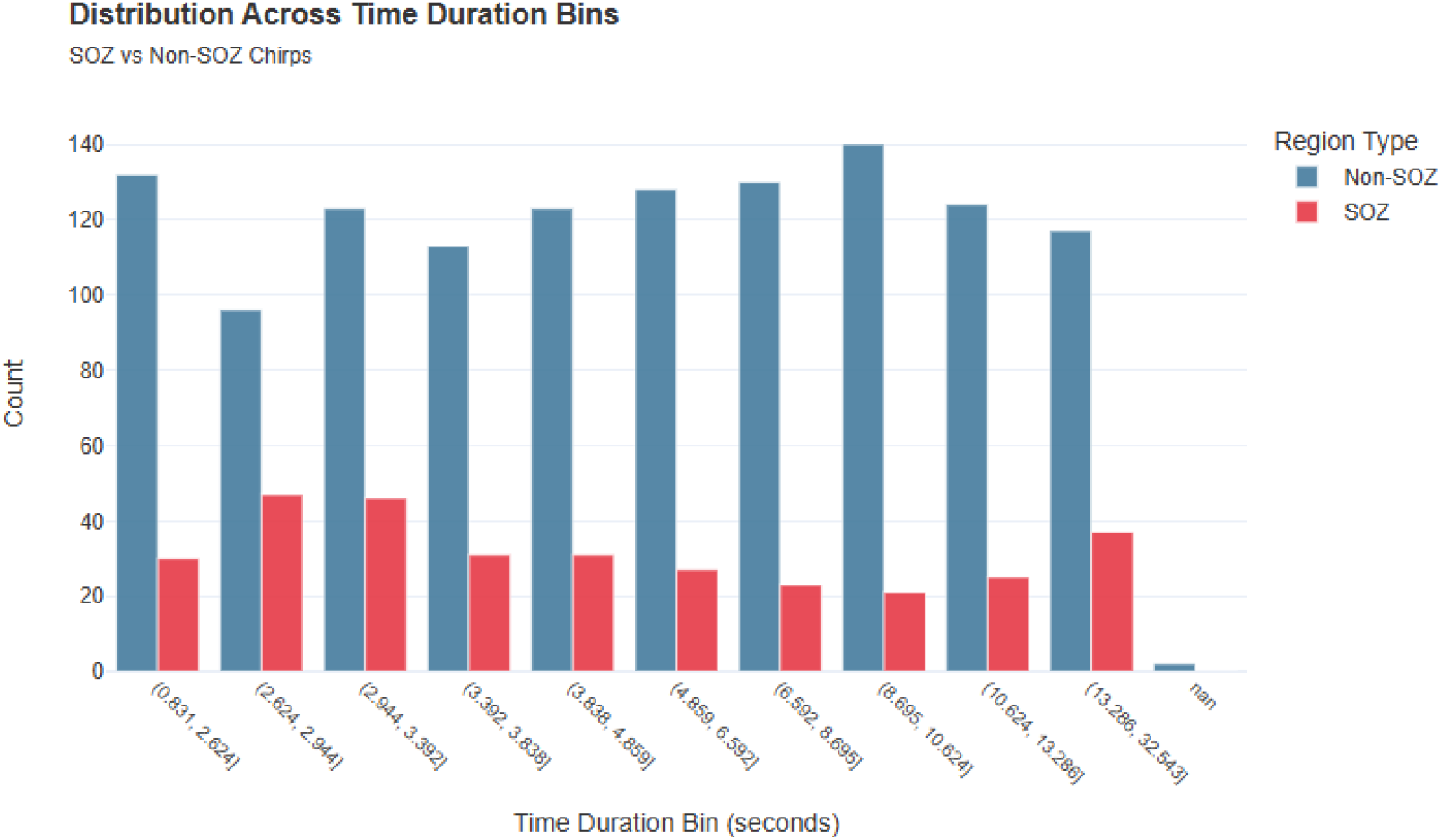
Distribution Across Temporal Duration Bins of CHIRPs: An opposing trend is observed between SOZ and Non-SOZ regions across increasing temporal duration bins. As the time duration of CHIRPs increases, from shorter to longer bins, the proportion of Non-SOZ CHIRPs (blue bars) generally rises, while the proportion of SOZ CHIRPs (red bars) steadily declines. This inverse relationship indicates that seizure onset zones tend to generate more short-duration CHIRPs, whereas non-seizure regions are associated with a higher frequency of long-duration CHIRPs. This pattern highlights a clear divergence in temporal characteristics of CHIRPs between epileptogenic and non-epileptogenic brain areas.

## 4. Discussion

The research conducted by Di Giacomo et al. (2024) provides evidence that fast activity “chirp” patterns observed in SEEG recordings can serve as dependable markers of the epileptogenic zone (EZ) in individuals with focal drug-resistant epilepsy. In this retrospective study of 176 patients, chirps—defined as brief, high-frequency signals that gradually shift in frequency—were consistently found in over 95% of seizure events involving low-voltage fast activity, regardless of underlying pathology or lobar origin. Crucially, the study found a significant association between chirp-EZ overlap and favorable surgical outcomes, with patients showing greater alignment between chirp sites and visually defined EZs more likely to achieve Engel class I results (p < .036). In contrast, discrepancies between chirp distribution and visually identified EZs were linked to poorer seizure control (p = .01). Among the subgroup of patients with complete seizure freedom (Engel class Ia), those who underwent resection or thermocoagulation of chirp-generating regions had particularly positive outcomes. The findings suggest that chirp-based spectral analysis offers a valuable, semi-automated complement to traditional EZ localization, especially in complex or MRI-negative cases, enhancing both diagnostic precision and surgical planning (Di Giacomo et al., 2024). While Di Giacomo (Di Giacomo et al., 2024) links chirps to EZs, our work explores their prognostic power, shifting focus from “where chirps occur” (EZ localization) to “how chirp features predict outcomes” (prognostics). Our study reveals several key findings about ictal chirps and their clinical relevance: Chirps were detected in both SOZ (20.6%) and non-SOZ (79.4%) regions, suggesting a broader network involvement beyond epileptogenic zones. However, SOZ chirps exhibited longer spectral durations (*p < 0*.*001*) and higher spectro-temporal ratios (*p < 0*.*001*), potentially reflecting more rapid frequency modulation during seizure initiation (Figure 10–12). Heatmap analysis (Figure 4) identified patient subgroups with similar chirp profiles (e.g., PT10/PT1, JH101/JH108), hinting at shared pathophysiological mechanisms. Hierarchical clustering of clinical profiles (Figures 5–6) grouped patients based on Engel/ILAE scores and geometric shape metrics, revealing distinct phenotypic clusters. Notably, patients with favorable outcomes (e.g., Engel class I) were enriched in clusters with shorter temporal chirp durations (Figure 14). Outcome prediction models (Figure 7) highlighted that SOZ chirps with very high temporal duration carried the highest risk of poor outcomes (51.4% risk), while non-SOZ chirps in medium spectral duration bins predicted worse outcomes (53.7% risk). Our semi-automated annotation pipeline (Figures 1–2) balanced human expertise with computational precision, enabling scalable chirp characterization.

## Limitations and Future Directions

The modest effect sizes (*r = 0.07–0.12*) suggest chirp features alone may be insufficient for definitive SOZ localization, necessitating integration with other biomarkers. Future studies could explore chirp dynamics in relation to seizure propagation networks or pharmacological response.

## Acknowledgments

The authors gratefully acknowledge the financial support of the Canadian Neuroanalytics Scholars Program and The Hilary & Galen Weston Foundation. We also recognize the valuable support provided by Campus Alberta Neuroscience, the Hotchkiss Brain Institute at the University of Calgary, the Ontario Brain Institute, and the Neuro at McGill University. Their contributions were essential to the advancement of this research project.

## Appendix I

Table 2 provides a summary of spatial, spectro-temporal, and clinical features analyzed in this study.

**Table 2.**
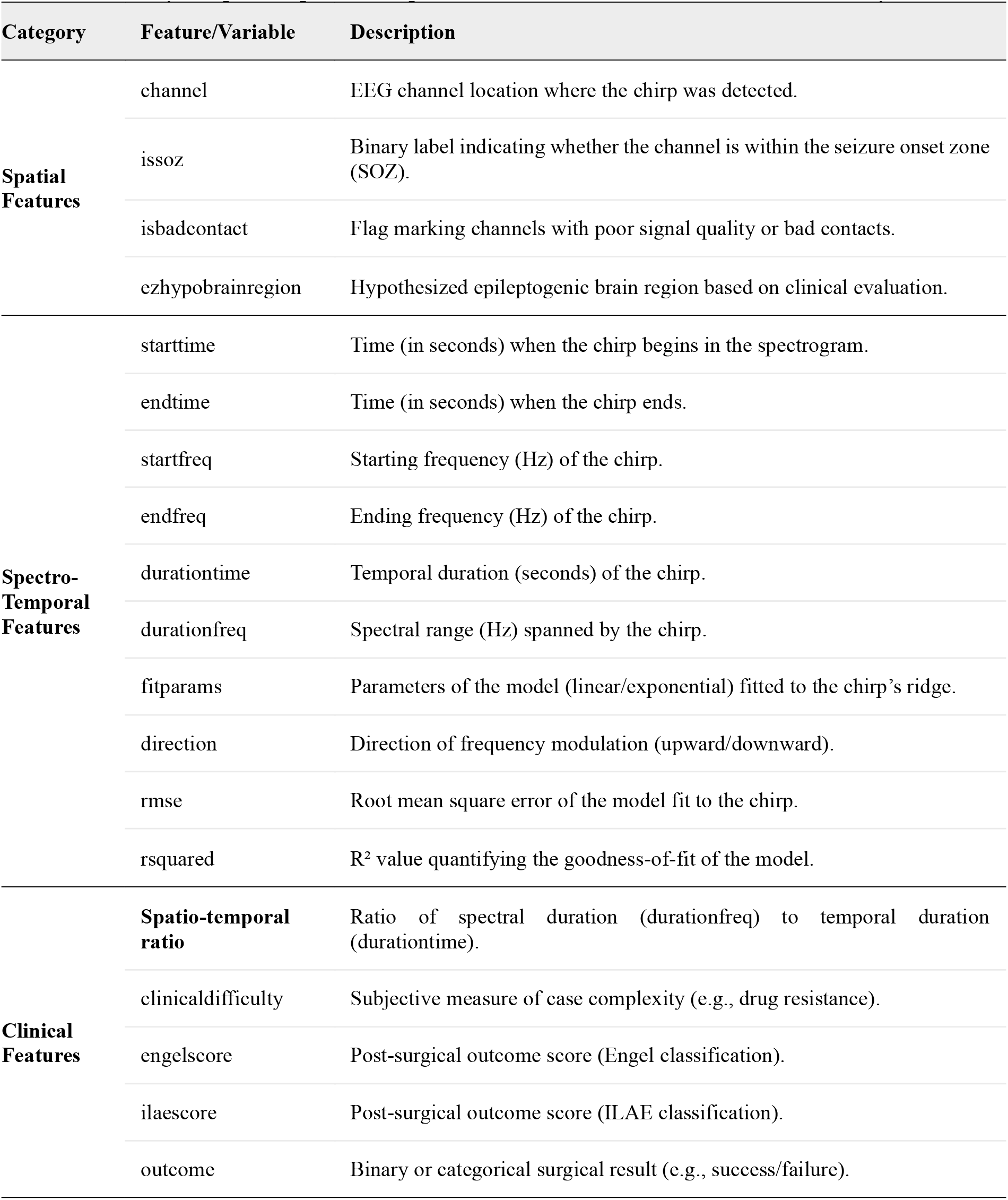
Summary of Spatial, Spectro-Temporal, and Clinical Features Evaluated in the Study.

## References

Adam Li and Sara Inati and Kareem Zaghloul and Nathan Crone and William Anderson and Emily Johnson and Iahn Cajigas and Damian Brusko and Jonathan Jagid and Angel Claudio and Andres Kanner and Jennifer Hopp and Stephanie Chen and Jennifer Haagensen and Sridevi Sarma (2023). Epilepsy-iEEG-Multicenter-Dataset. OpenNeuro. [Dataset] doi: doi:10.18112/openneuro.ds003029.v1.0.6.

Bahador, N., Lankarany, M. (2025). Chirp Localization via Fine-Tuned Transformer Model: A Proof-of-Concept Study. arXiv preprint 2503.22713.

Bahador, N., Skinner, F., Zhang, L., & Lankarany, M. (2024). Ictal-related chirp as a biomarker for monitoring seizure progression. doi:10.1101/2024.10.29.620811.

Benedetto, J. J. and Colella, D. (1995). Wavelet analysis of spectrogram seizure chirps. In Wavelet Applications in Signal and Image Processing III, volume 2569, pages 512–521. SPIE.

de Curtis, M. and Avoli, M. (2016). GABAergic networks jump-start focal seizures. Epilepsia, 57(5):679–687.

Di Giacomo, R., Burini, A., Chiarello, D., Pelliccia, V., Deleo, F., Garbelli, R., De Curtis, M., Tassi, L., & Gnatkovsky, V. (2024). Ictal fast activity chirps as markers of the epileptogenic zone. Epilepsia, 65(6). 10.1111/epi.17995.

Feltane, A., Bartels, G. B., Boudria, Y., and Besio, W. (2013). Analyzing the presence of chirp signals in the electroencephalogram during seizure using the reassignment time-frequency representation and the hough transform. In 2013 6th International IEEE/EMBS Conference on Neural Engineering (NER), pages 186–189. IEEE.

Freeman, J. M., Vining, E. P., and J., P. D. (1993). Seizures and epilepsy in childhood: a guide for parents.

Frosz, M. H. and Andersen, P. E. (2007). Can pulse broadening be stopped? Nature Photonics, 1(11):611–612.

Gnatkovsky, V., Francione, S., Cardinale, F., Mai, R., Tassi, L., Lo Russo, G., and De Curtis, M. (2011). Identification of reproducible ictal patterns based on quantified frequency analysis of intracranial eeg signals. Epilepsia, 52(3):477–488.

Gnatkovsky, V., Pelliccia, V., de Curtis, M., and Tassi, L. (2019a). Two main focal seizure patterns revealed by intracerebral electroencephalographic biomarker analysis. Epilepsia, 60(1):96–106.

Grinenko, O., Li, J., Mosher, J. C., Wang, I. Z., Bulacio, J. C., Gonzalez-Martinez, J., Nair, D., Najm, I., Leahy, R. M., and Chauvel, P. (2018). A fingerprint of the epileptogenic zone in human epilepsies. Brain, 141(1):117–131.

Kurbatova, P., Wendling, F., Kaminska, A., Rosati, A., Nabbout, R., Guerrini, R., Dulac, O., Pons, G., Cornu, C., Nony, P., et al. (2016). Dynamic changes of depolarizing gaba in a computational model of epileptogenic brain: Insight for dravet syndrome. Experimental Neurology, 283:57–72.

Li, J., Grinenko, O., Mosher, J. C., Gonzalez-Martinez, J., Leahy, R. M., and Chauvel, P. (2020). Learning to define an electrical biomarker of the epileptogenic zone. Human Brain Mapping, 41(2):429–441.

Miri, M. L., Vinck, M., Pant, R., and Cardin, J. A. (2018). Altered hippocampal interneuron activity precedes ictal onset. eLife, 7:e40750.

Niederhauser, J. J., Esteller, R., Echauz, J., Vachtsevanos, G., and Litt, B. (2003). Detection of seizure precursors from depth-eeg using a sign periodogram transform. IEEE Transactions on Biomedical Engineering, 50(4):449–458.

Rich, S., Chameh, H. M., Rafiee, M., Ferguson, K., Skinner, F. K., and Valiante, T. A. (2020). Inhibitory Network Bistability Explains Increased Interneuronal Activity Prior to Seizure Onset. Frontiers in Neural Circuits, 13.

Schiff, S. J., Colella, D., Jacyna, G. M., Hughes, E., Creekmore, J. W., Marshall, A., Bozek-Kuzmicki, M., Benke, G., Gaillard, W. D., Conry, J., et al. (2000). Brain chirps: spectrographic signatures of epileptic seizures. Clinical Neurophysiology, 111(6):953–958.

Sen, A., Kubek, M., and Shannon, H. (2007). Analysis of seizure eeg in kindled epileptic rats. Computational and Mathematical Methods in Medicine, 8(4):225–234.

Wieser, H. G., Blume, W. T., Fish, D., Goldensohn, E., Hufnagel, A., King, D., … Lüders, H. (2001). Proposal for a new classification of outcome with respect to epileptic seizures following epilepsy surgery. Epilepsia, 42(2), 282–286. doi:10.1046/j.1528-1157.2001.4220282.x.

